# Global distribution of cultivated *Trichodesmium* subclades revealed by multi-omics

**DOI:** 10.64898/2026.07.10.737770

**Authors:** Yiming Zhao, Catie S. Cleveland, Max R. Binkowski, Brianna Batson, Asa E. Conover, Eric A. Webb

**Author notes:** Corresponding Author: Eric A. Webb.

## Abstract

*Trichodesmium* is an important oceanic N_2_ fixing cyanobacterial genus that has been shown to provide up to 50% of new N to oligotrophic regimes. Despite its importance, we know relatively little about the genomic potential and functional diversity of the two major clades of *Trichodesmium* (Clade I and Clade III, hereafter Thieb and Tery, respectively). With the expanded collection of *Trichodesmium* isolates in the USCTCC (University of Southern California *Trichodesmium* Culture Collection), we sequenced genomes from seven cultivated strains to further characterize the genomic diversity within the genus. For example, sequencing the genome of the “gigantic”, red *Trichodesmium contortum* surprisingly shows that they are closely related to the smallest *Trichodesmium* clade, TeryA. The high genomic identity between TeryA and *T. contortum* (>98.5% ANI) and lack of implicated auxiliary genes suggests their large biovolume differences might be transcriptional or epigenetic in origin. Furthermore, these data show that using Tery-subclades are a more accurate designation than the classical *Trichodesmium* species delineation. Finally, we placed the analysis of these genomes in an ecological context via read mapping with globally distributed RNA and DNA datasets. Our data show that Tery clades (A&B) are both lower in relative abundance compared to Thieb in global oceans, generally co-occur when detected in the field, and are highly linked to decreased salinity and increased temperatures, especially for sampling locations in the Bay of Bengal. Lastly, even though TeryA members are undersaturated with respect to current CO_2_ concentrations, our phenotypic and biogeography data suggests that salinity/ocean color could limit their predicted global impact as climate changes.

## Introduction

Fixation of atmospheric N_2_ to NH_3_ in the oligotrophic ocean is recognized as an important source of new, bioavailable nitrogen (Sohm et al., 2011; Zehr & Capone, 2020). Generation of new nitrogen in the surface oceans can significantly influence primary production and the amount of organic material being exported through the biological pump (Longhurst & Glen Harrison, 1989; Sohm et al., 2011). Thus, revealing the diversity, abundances, and activity of N_2_-fixing organisms in the surface oceans is crucial to understand marine C and N cycling (Latysheva et al., 2012; Luo et al., 2012; Zehr, 2011).

The N_2_-fixing genus *Trichodesmium* has been widely acknowledged for significantly contributing to new N generation in tropical and subtropical oligotrophic ocean waters (Bergman et al., 2013; Capone et al., 1997; Letelier & Karl, 1996), where it can seasonally contribute approximately half of the microbially-derived N input regionally in subtropical gyres. Despite the prominent role of *Trichodesmium* in ocean biogeochemical cycling (Carpenter et al., 2004; Deutsch et al., 2001; Martin et al., 2019; Montoya et al., 2004; Sohm et al., 2011; Zehr & Capone, 2020), its evolutionary history, clade-defining characteristics, and *in situ* distribution of the genus remain unclear.

Morphological studies historically defined six species of *Trichodesmium* (*T. thiebautii, T. tenue, T. pelagicum, T. hildebrandtii, T. erythraeum,* and *T. contortum*) (Bergman et al., 2013; Hynes et al., 2012; Lundgren et al., 2001; Webb et al., 2022). Genetic analysis of recent laboratory isolates and environmental metagenomic assemblies have since shown two major diazotrophic clades of *Trichodesmium* (*T. erythraeum* or Tery, and *T. thiebautii* or Thieb) composed by four total subclades (Tery-A&B, Thieb-A&B (Webb et al., 2023)). But within the Tery clade specifically, there were previously only three genomes from cultivated strains representing TeryA (i.e., *T. erythraeum* strains IMS101, 2175, and LIN; (Walworth et al., 2015; Conover et al., 2021; Webb et al., 2023)), while representatives of TeryB went extinct in culture collections before their genomes could be sequenced (Hynes et al., 2012). There is also only one whole genome from a cultivated strain representing Thieb (strain H94; (Webb et al., 2023)). Of these few cultivated *Trichodesmium* strains, Tery have been characterized as faster growing than Thieb, having a higher thermal optimum (Chappell & Webb, 2010), and less CO_2_-saturated (Hutchins et al., 2013; Gradoville et al., 2014). Based on these data, it was hypothesized that Tery would be more successful under climate change and thus have an increased role in biogeochemical cycling, C+N export, and community structure in future oceans.

In this study, we performed a multi-omic analysis of the genetic potential of the different *Trichodesmium* clades to respond to their environment. To do so, we generated high quality genomes from 7 previously unsequenced Tery isolates (2 TeryA, 5 TeryB). Our work shows that despite Tery’s generally faster growth and CO_2_ undersaturation, Thieb numerically dominates the current oceans in metagenomic samples with corresponding high activity in RNAseq datasets. These results challenge our view of Tery contribution in global N cycling and call for a closer look at *Trichodesmium* biogeochemical contribution at the subclade level.

## Material and Methods

### *Trichodesmium* isolates maintenance

New *Trichodesmium* isolates were either obtained from John Waterbury at the Woods Hole Oceanographic Institution (strains GS4 and St8 isolated in 75% seawater-based RMP or PMP; Webb et al 2001; Hynes et al 2012), or isolated by members of the Webb Lab from the Trans-Atlantic TriCoLim cruise or the Bermuda Atlantic Time-series Station (BATS) using either RMP or YBCII (Chen et al 1996). The enrichments were set up shipboard from hand-picked colonies as described in Webb et al., 2022. USCTCC isolates used in this work were maintained in artificial seawater media YBCII (Chen et al., 1996) and kept with 50-150 µEi m^-1^s^-1^ warm-white fluorescent light on a 12:12 light:dark diel cycle at ∼26° C in Percival incubators (Perry, Iowa, USA).

### Microscopic visualization and cell size measurement

*Trichodesmium* lab isolates were visualized, photographed, and cell sizes were measured using an AxioStar Plus microscope (Zeiss, Okerkochen, Germany) with X-Cite lamp (Excelitas, Pittsburgh, PA, USA) and DAPI Long Pass filter. To visualize filament color change, light exposure of *Trichodesmium* isolates experiments were done with the same lamp and filter but with longer exposure times (1-20mins).

### DNA extraction and sequencing

Biomass for DNA extraction was collected from ∼100ml dense cultures via gentle vacuum filtration onto a 25mm 5-µm polycarbonate filter (Whatman, Piscataway, New Jersey, USA) for all strains excluding *T. erythraeum* CON. The latter was filtered onto 40-µm cell strainers (Corning, Glendale, Arizona, USA) to avoid the filter clogging from the high amounts of transparent exopolysaccharides (TEP) this isolate produces. DNA extractions of *T. erythraeum* RLI, *T. erythraeum* DEMO, and *T. erythraeum* GS4 were done using Qiagen DNeasy Powersoil Kit (Germantown, MD) according to manufacturer’s protocol. DNA extractions of *T. erythraeum* CON, *T. erythraeum* ST8, *T. erythraeum* T5, *T. erythraeum* T114 were done with the Phenol-chloroform-CTAB protocol (Urakawa et al., 2010). DNA was validated as high-quality before sequencing using NanoDrop UV-Vis spectrophotometer (ThermoFisher, Waltham, MA, USA) and Qubit dsDNA Quantification Assay Kits (ThermoFisher, Waltham, MA, USA). The DNA samples were then further evaluated and processed by Novogene (Sacramento, CA, USA) using the NEBNext DNA library Prep Kit and a 150 Illumina PE sequencing protocol.

### Genome assembly

All seven new isolate assemblies (*T. erythraeum* RLI, *T. erythraeum* CON, *T. erythraeum* DEMO, *T. erythraeum* ST8, *T. erythraeum* GS4, *T. erythraeum* T5, and *T. erythraeum* T114) were processed on the University of Southern California Center for Advanced Research Computing (USC CARC) via the pipeline briefly described below. Raw reads quality were first checked with FastQC v0.12.1 (Andrews, 2010) and then trimmed with Trimmomatic v0.39 (Bolger et al., 2014). Both MEGAHIT v1.2.9 (Li et al., 2015) and metaSpades v3.13.0 (Nurk et al., 2017) were used to assemble genomes. Assemblies from both assemblers essentially showed the same stats (i.e., #contigs and N50), thus assemblies from metaSpades were kept for the downstream steps. Then assemblies were binned by MaxBin2 v2.2.4 (Wu et al., 2016) and further screened with CheckM v1.0.18 (Parks et al., 2015) to determine completeness and contamination.

### Genome refining

Fourteen genomes (including: 7 new isolate assemblies, 4 previously published isolate genomes, and 3 previously published MAGs (**Supp. Table 1**)) were refined based on Kaiju v.1.10.1 contig taxonomy classification with the NCBI nr database and GC percentage within Anvi’o v.8 interactive interface. Here, contigs were removed if they had not been taxonomically assigned as Cyanobacteria in Kaiju and displayed significantly higher/lower GC content than *Trichodesmium* sequences in the same genome.

### Phylogenomics

All genomes from isolates used in this study were run through GToTree v1.7.06 (Lee, 2019) using 251 cyanobacterial core proteins Hidden Markov Models (HMMs) to determine initial phylogenetic placement with *Synechocystis* sp. PCC 6803 (GCA_000340785.1) as the root of this maximum likelihood (ML) phylogeny (FastTree2; Price et al., 2010). The output files generated from GToTree were fed into IQtree v2.0.3 (Nguyen et al., 2015) in ModelFinder optimality mode and 1000 resampling rounds to improve tree models. The final ML tree was visualized in the iTol web server (Letunic & Bork, 2021) and edited in Affinity Designer (Nottingham, United Kingdom).

### Pigmentation extraction and absorption spectra analysis

To optimize the production of pigments, cultures were acclimated to low light (∼30 µEi m^-1^s^-1^) for at least 1 month and at least 2 transfers. Pigments were extracted similar to previous methods (Bell & Fu, 2005; Hynes et al., 2012) with the following specifications: ∼200 mL dense culture was filtered down onto 5-µm polycarbonate filters (Whatman, Piscataway, New Jersey, USA), filters were frozen short term at -20°C, thawed and re-suspended off filters in 4°C 1x phosphate-buffered saline (PBS), sonicated for 30 sec in 1s on/off intervals with amplitude of 0.6. These extracts were then ultracentrifuged (Beckman Coulter, San Jose, California, USA) for 1 hour at 4°C at 100,000 rpm (>200,000 x g), and the 400-750 nm absorbance spectra was measured in 10 nm steps on the SpectraMAX M2e (San Jose, California, USA) plate reader in Quartz cuvettes. A PBS blank was subtracted from both spectra, and they were normalized to the largest peak absorbance (max. at 490 nm for DEMO).

### Pangenomics

Pangenomic analysis was performed with the Anvi’o v7.1 pangenome pipeline (Eren et al., 2015, 2021) using all 14 genomes mentioned above. The single copy *Trichodesmium* core genetic content identified in this step was also used as a read-mapping reference database in metagenomic pipelines. Briefly, contig databases were created for each genome using Anvi’o contigs database generating function (which automatically computed k-mer frequencies with default of 4), open reading frames were identified with Prodigal v2.6.3 (Hyatt et al., 2010), and visualized on an interactive page prompted in Anvi’o (Eren et al., 2015, 2021). It is important to note that we did not consider singletons (gene clusters that are only present in one single genome) in our downstream metagenomic analysis, since almost all genomes, although high quality, are not closed.

### Metagenomics and global map

Metagenomic sequences of global bulk water samples from GO-SHIP, bioGEOTRACES, station ALOHA and BATS (Biller et al., 2018; Larkin et al., 2021) were used for read-recruiting to understand *Trichodesmium* biogeography. Due to the highly similar nucleotide content between subclades (i.e., >98.5% between TeryA-1&2, and also >98.5% between TeryB-1&2), there was a higher risk of cross-mapping during metagenomic read recruitment, thus we only use core genome in *T. erythraeum* IMS101 to represent TeryA and core genome in *T. erythraeum* DEMO to represent TeryB; we could not further resolve the TeryAB-1&2 subclades based on core genome similarity alone.

Raw reads from bulk water samples mentioned above were recruited to this contig database using bowtie2 v2.4.1 (Langmead & Salzberg, 2012), which outputs total read-recruitment percentage of *Trichodesmium* at each sampling location. Samtools v18.0.4 (Danecek et al., 2021) converted the Sequence Alignment Map (SAMs) files into Binary Alignment Maps (BAMs), and CoverM v0.4.0 (Aroney et al., 2025) was then used with an identity threshold of 0.98 to filter potential cross-read-recruitment due to the highly similar genomes in the reference contig database or errant bowtie2 mapping with default parameters. After filtering, the indexed BAMs were processed in Anvi’o v7.1 to determine relative abundance of different representative strains at each sampling station. Both total *Trichodesmium* read-recruitment percentage and relative abundance of each strain were used to generate the global distribution figure with the R package ggplot2 v3.4.2 (Villanueva & Chen, 2019). Metagenomes sequences from picked *Trichodesmium* colonies from the 2018 trans-Atlantic Ocean TriCoLim cruise (Webb et al., 2023) were also used to understand colony composition and distribution across the North-Equatorial Atlantic region.

### Metatranscriptome read recruiting

*Trichodesmium* colony metatranscriptomes from previous field studies (Cerdan-Garcia et al., 2022; Frischkorn et al., 2018; Rouco et al., 2018) were downloaded from NCBI and read-recruited to the same single copy core contig database to estimate the “activity” of each subclade in situ. Read-recruitment results from these datasets were then merged and visualized using Anvi’o and R for reads from Atlantic and Pacific oceanographic regions.

## Results and Discussion

### Phylogenomic placement and morphological characteristics of *Trichodesmium* isolates

#### a. Four subclades found inside *T. erythraeum* (the ‘Tery’ clade)

Phylogenomic analysis of all isolate genomes used here shows clear, well-supported branching pattern representing subclades TeryA-1, TeryA-2, TeryB-1, and TeryB-2 in the previously defined ‘Tery’ clade (Webb et al., 2023) (**Fig. 1**). *T. tenue/thiebautii* H94 sequenced by Webb et al. (2023) is also included to highlight the species-level phylogenomic differences. H94 has been previously phylogenetically placed using marker genes in a clade with *T. tenue*, *T. spiralis*, and *T. hildebrandtii* (Hynes et al., 2012), and studies have defined it as both *T. tenue* and *T. thiebautii* (Chappell & Webb, 2010; Hynes et al., 2012; Walworth et al., 2015). In this study, we will refer to this as ‘Thieb’ (representing *T. thiebautii*) based on the studies with genetic analysis (Hynes et al., 2012; Webb et al., 2023).

**Figure 1.**
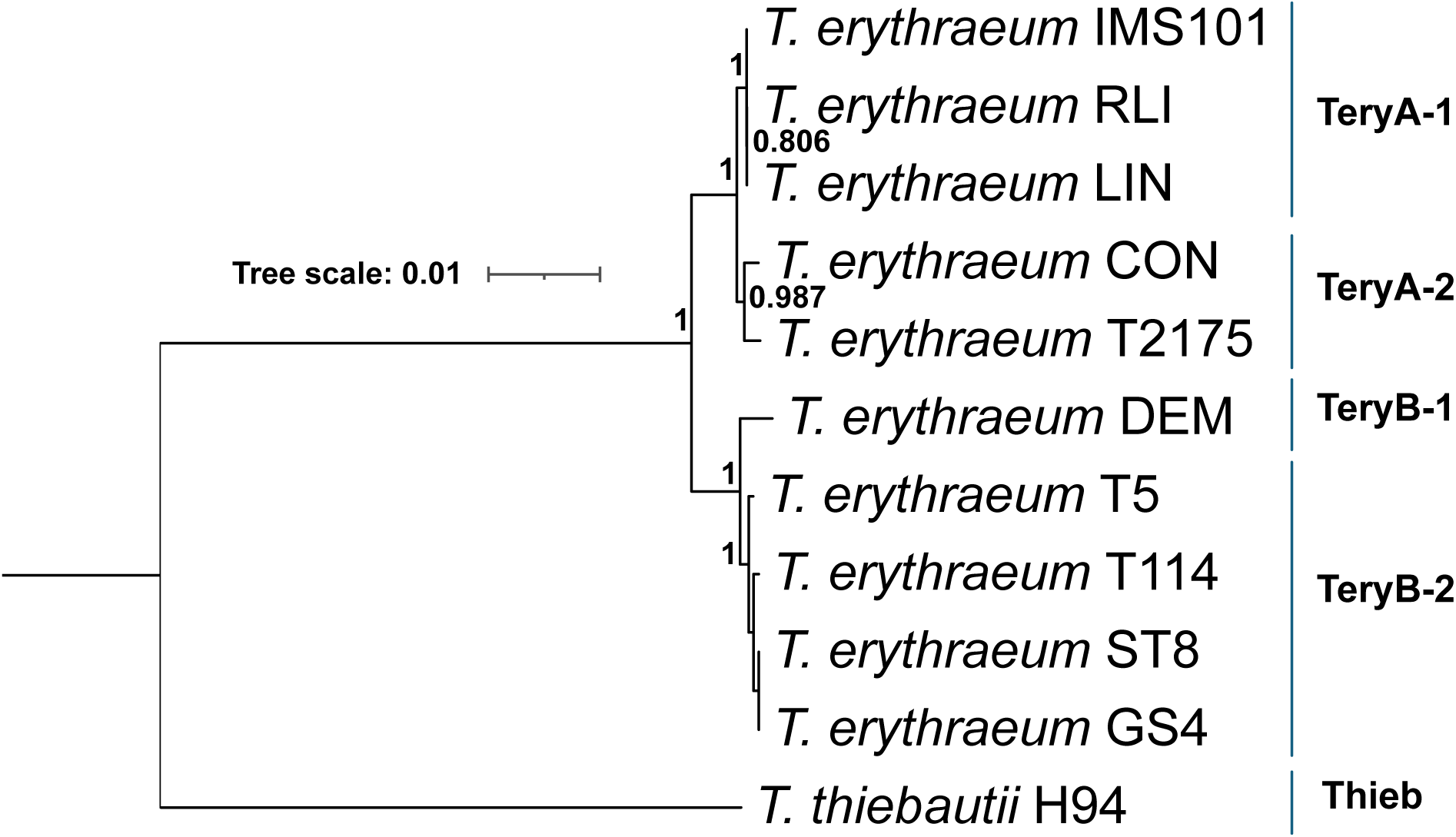
Phylogenomic tree of *Trichodesmium* isolate genomes with subclade assignments on the right. The genomes included in phylogenomic tree represent 4 subclades (TeryA-1, TeryA-2, TeryB-1, TeryB-2) within *T. erythraeum*, and one genome representing *T. thiebautii* (Thieb). The tree is rooted with *Synechocystis sp.* PCC6803 (not shown), and only bootstrap values above 0.5 are shown.

Lab-grown phenotypes are broadly consistent with this phylogenomic placement. For example, both TeryA-1 and TeryA-2 do not form colonies during exponential lab growth, and all cultures show bright red color when visualized with a DAPI long-pass filter (**Fig. 2A-D, Table 1**). These phenotypes were also observed in other extinct isolates in Hynes et al., (2012) and discussed in Webb et al., (2023). Conversely, all TeryB-1, TeryB-2, and Thieb representatives often form colonies in “tufts” and are red to yellow-green ((Hynes et al., 2012); **Fig. 2E-F**, **Table 1**). In general, these data showed that the phylogenomics generally tracks with observable phenotypic differences in the subclades.

**Figure 2.**
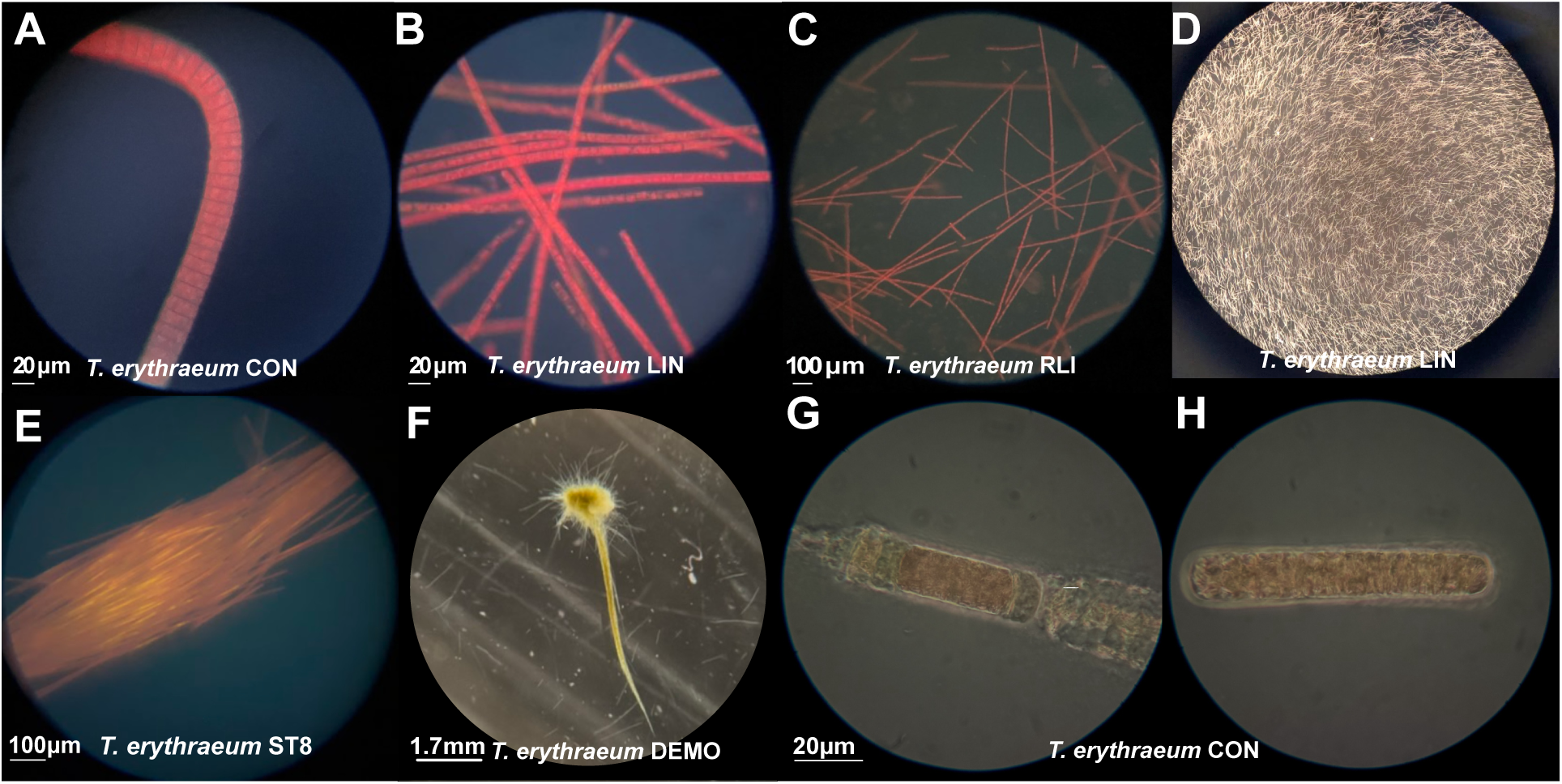
General morphology of various *Trichodesmium* strains. (**A-C**) Cell size comparison between *T. erythraeum* CON (TeryA-2), *T. erythraeum* LIN&RLI (TeryA-1). Photographs were taken using DAPI-longpass filter, illustrating the distinctive red epifluorescent color of *T. erythraeum* cells. (**D**) Dissecting microscopic image of *T. erythraeum* LIN illustrating culture density and absence of colony formation. (**E)** *T. erythraeum* ST8 (TeryB-2) under near UV light (**F**) *T. erythraeum* DEMO under incandescent light suggests unique green color. (**G**) Cellular lysis of *T. erythraeum* CON and the formation of a smaller filament (i.e., perhaps hormogonium-like structure), and (**H**) the short structure of *T. erythraeum* CON cell-stacks that survived lysis of longer trichome. In **H**, the hormogonium-like structure from a stressed *T. erythraeum* CON culture was stable for >2weeks. Interestingly, clean cellular edges can be seen on the trichome ends (as opposed to just-lysed trichome ends in **G**).

**Table 1.**
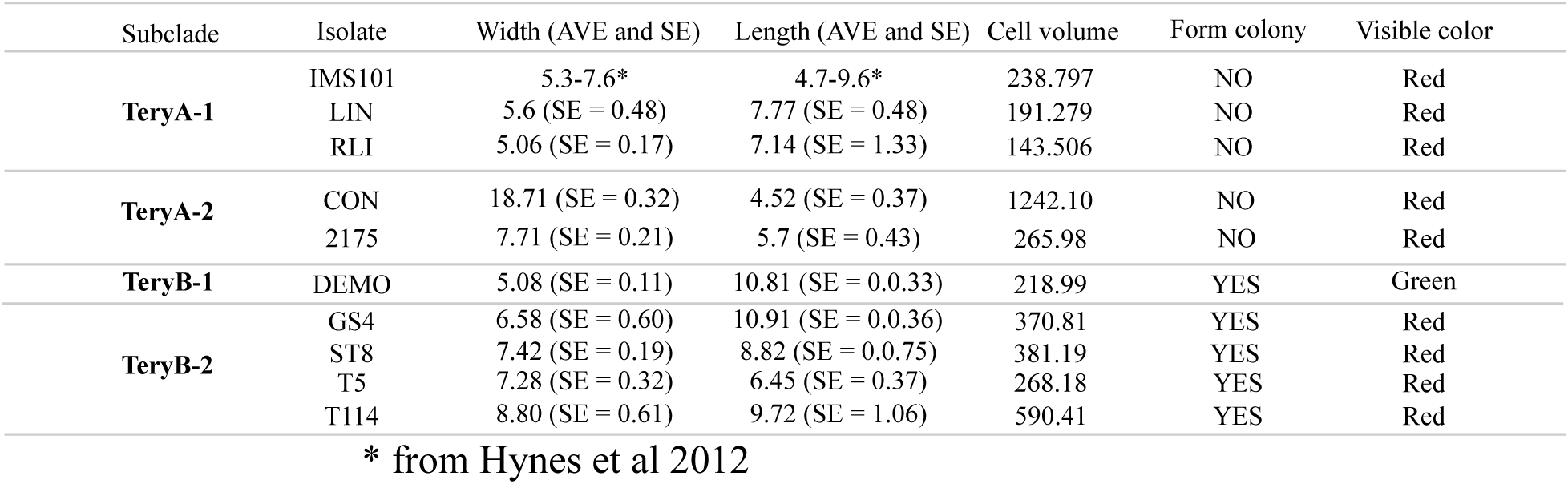
Cell size and phenotypic features of *T. erythraeum* subclades. Width and length unit: µm.

Although phylogenomic placement showed TeryA-1 and TeryA-2 as very closely related, their cell characteristics can be quite different (**Table 1**). We found that TeryA-2 cells are wider than their length and have disk-like cell shape, while TeryA-1 cells are longer than their width (**Table 1** & **Fig. 2**; (Hynes et al., 2012; Janson et al., 1995)). Strains within TeryA-2 are much more variable than TeryA-1 in cell volume. While the cell volume of TeryA-2 strain 2175 is not significantly different than TeryA-1, cells of TeryA-2 CON are >5 times larger on average.

We hypothesize that the ‘wider than longer’ cell shape of TeryA-2 indicates potentially higher needs of inter-cell communication than cell-environment exchange, as the wider cell shape gives a much larger contacting surface between two connected cells inside a trichome. This finding also has implications for their external environment-exposed surface area, likely making diffusion less favorable in TeryA-2 relative to TeryA-1, which has a larger exposed area. Our hypotheses would need further molecular evidence, and more TeryA-2 strains are needed in lab culture collections to attempt the identification of subclade-specific genetic content responsible for this feature. Overall, differences in intracellular biovolume and C+N content could lead to differences in TeryA-2 contribution to C+N standing stocks and biogeochemical cycling and thus should not be overlooked when discussing *Trichodesmium* nutrient cycling contributions.

Some strains have characteristics not found among others, either within their sub-clade or more broadly. We found that TeryB-1 DEMO appears to be optimized to absorb blue light, while other Tery strains were found to absorb mostly green light (Hynes et al. 2012). Specifically, spectral analyses showed that TeryB-1 DEMO is the only Tery strain to show a blue-shifted profile of phycourobilin (PUB, 495nm): phycoerythrobilin (PEB, 545nm) with a pigment ratio value of 1.62. Previously, blue-tuned PUB:PEB ratios (>1) were only demonstrated in Thieb strains (**Fig. 2F**, (Hynes et al., 2012).

#### b. TeryA cladistic and phenotypic correlation

Previous studies have put in efforts determining *Trichodesmium* cladistics using both metagenomic assembled genomes (MAGs) and marker genes (Hynes et al., 2012; Janson et al., 1999; Koedooder et al., 2022; Webb et al., 2023). However, there are disadvantages of using only one of these two paths. The high average nucleotide identity (ANI) and large portion of non-coding DNA among *Trichodesmium* genomes (Walworth et al., 2015; Webb et al., 2023) can cause assembly artifacts in mixed samples, while marker gene-based phylogenetics have worse resolution and can be misleading (Koedooder et al., 2022). The latter point is important to consider especially during the analysis of MAGs.

For example, a recent study in the Red Sea noted that using *hetR* phylogenies to determine *Trichodesmium* diversity can lead to overestimations and confusion in terms of *Trichodesmium* cladistics because some *Trichodesmium* genomes have two copies of *hetR* (Koedooder et al., 2022). Indeed, our newer Thieb H94 genome has two phylogenetically disparate *hetR* genes (data not shown). However in addition to this, Koedooder et al used both the ANI of MAG R03 that clustered with non-diazotrophic *T. nobis* and the high identity of its *hetR* sequence to a previously designated *T. contortum* colony (Janson et al., 1999) to conclude that *T. contortum* should be a close relative to non-diazotrophic *Trichodesmium* (Koedooder et al., 2022). Contrastingly, our combined phylogenomic and phenotypic data indicated that TeryA subclade TeryA-2 (previously defined *T. contortum* (Janson et al., 1994)) and TeryA-1 (previously defined *T. erythraeum* (Gomont, 1893)) are very closely related **(Fig. 1&2)** and relatively “distant” from the non-diazotrophic *Trichodesmium*.

One possible explanation of mismatching results between our work and others (Janson et al., 1994; Koedooder et al., 2022), could be due to extraction bias during sample preparation. We noticed TeryA-2 CON produces visibly excessive amounts of transparent high viscosity material around the trichomes, which leads to the abnormal difficulty of extracting DNA from this strain. Thus, for the original Janson et al (Janson et al., 1999) “*T. contortum”* samples, we hypothesize that there could have been some fixed-nitrogen utilizing, non-diazotrophic *Trichodesmium* co-existing in Janson’s *T. contortum* sample that got extracted, *hetR* amplified, cloned, and sequenced instead. Thus, when these ‘*T. contortum’ hetR* sequences from Janson’s work were used in the Red Sea study (Koedooder et al., 2022), they were erroneously attributed to *T. contortum* but had in fact originated from “cheating” non-diazotrophic *Trichodesmium*.

Our argument agrees with the close *hetR* placement with *T. erythraeum* isolates and disc-shape phenotypes of extinct *T. contortum* strains 21-74 and 20-70 described by Hynes (Hynes et al 2012). Contrarily, in the absence of non-diazotrophic *Trichodesmium* cultures, it is possible that one or both of them are phenotypically similar to *T. contortum.* However as discussed above this disc-shaped morphology would make the non-diazotrophs less efficient at diffusion, a likely non-successful phenotype for the oligotrophic oceans. Importantly, the phenotype and the genotype of our TeryA-2 strain CON, emphasizes the importance of using whole genomes of *Trichodesmium* isolates for phylogenetic placement, agrees with previous marker gene phylogenetics of extinct cultures (Hynes et al., 2012), and clearly shows that the TeryA-2 clade has larger, red disc-shaped cells.

### *T. erythraeum* CON showed the first possible evidence of hormogonia-like cells in *Trichodesmium*

Filamentous cyanobacteria, including many members of the *Oscillatoriales* which includes *Trichodesmium*, are known to make smaller differentiated cells that function in dispersal and phototaxis called hormogonia (Risser, 2023). While hormogonia have never been documented in *Trichodesmium*, *T. erythraeum* CON (TeryA-2) was visualized with potential hormogonium production under strong light exposure from a microscope and/or under nutrient stress in lab conditions. When grown with replete nutrients and light levels around 100 µEi m^-1^s^-1^, CON trichomes are elongated in length and can bend due to the long trichome formed with disc-shape cells (**Fig. 2A**). Under high light stress from the microscope, the CON trichomes started a ‘chain-reaction’ of cellular lysis from the trichome poles one by one until only a few cells would remain (**Fig. 2G**). The ‘surviving’ region can be 5-20 cells long and stable under the same light stress for more than 20mins. Similarly, CON stationary cultures exhibited this same shortened trichome phenotype (**Fig. 2H**). These visualizations are consistent with the description of hormogonia in previous studies with *Nostoc spp*. (Adams & Duggan, 2008; Risser, 2023), where they demonstrated hormogonium production benefits the buoyancy and motility of such cells. Although motile behavior was not observed in *Trichodesmium* subclades, gliding motility is common in the genus (Pfreundt et al., 2023). Additionally, we do note that the CON culture was isolated from a likely Thieb colony (based on pigmentation) without any observable TeryA-2 characteristics present in that original field-picked colony (e.g., pigmentation or cell size). Thus, it is possible that the cellular differentiation facilitates CON cells leaving a dying *Trichodesmium* colony via increased buoyancy of these hormogonia-like CON cells to the surface and dispersing to a new colony. Further studies are needed to test the motility and phototaxis of these hormogonia-like cells to verify the function of this morphology in *Trichodesmium*.

### Tery subclades showed varied real-time color change phenotype under light stress

We observed that all Tery subclades can change color in real-time when exposed to strong, blue-shifted light (∼365nm) wavelength and visualized with a DAPI-longpass filter on an epifluorescent microscope (**Fig. 3**). Interestingly, not all subclades displayed the same level or pattern of color change. For example, TeryB-1 DEMO had the quickest color change in an expanding pattern (**Fig. 3A, supplemental material video**). Based on numerous observations over more than three years, once the blue light hit DEMO biomass, it started glowing golden in the center of the light field for ∼30s, then the glowing center expanded quickly and became about three times bigger within one minute. The expansion slowed at the light field edge, indicating the color change is likely in response to light shock. TeryB-2 isolates also changed to a similar golden color, but not as fast (**Fig. 3B**) - it can take 5-20mins for TeryB-2 members to significantly change color. Additionally, TeryB-2 colonies’ color didn’t change in the expanding manner as shown in TeryB-1 DEMO. Instead, random trichomes would change color and turn the colony into a speckled pattern (**Fig. 3B**).

**Figure 3.**
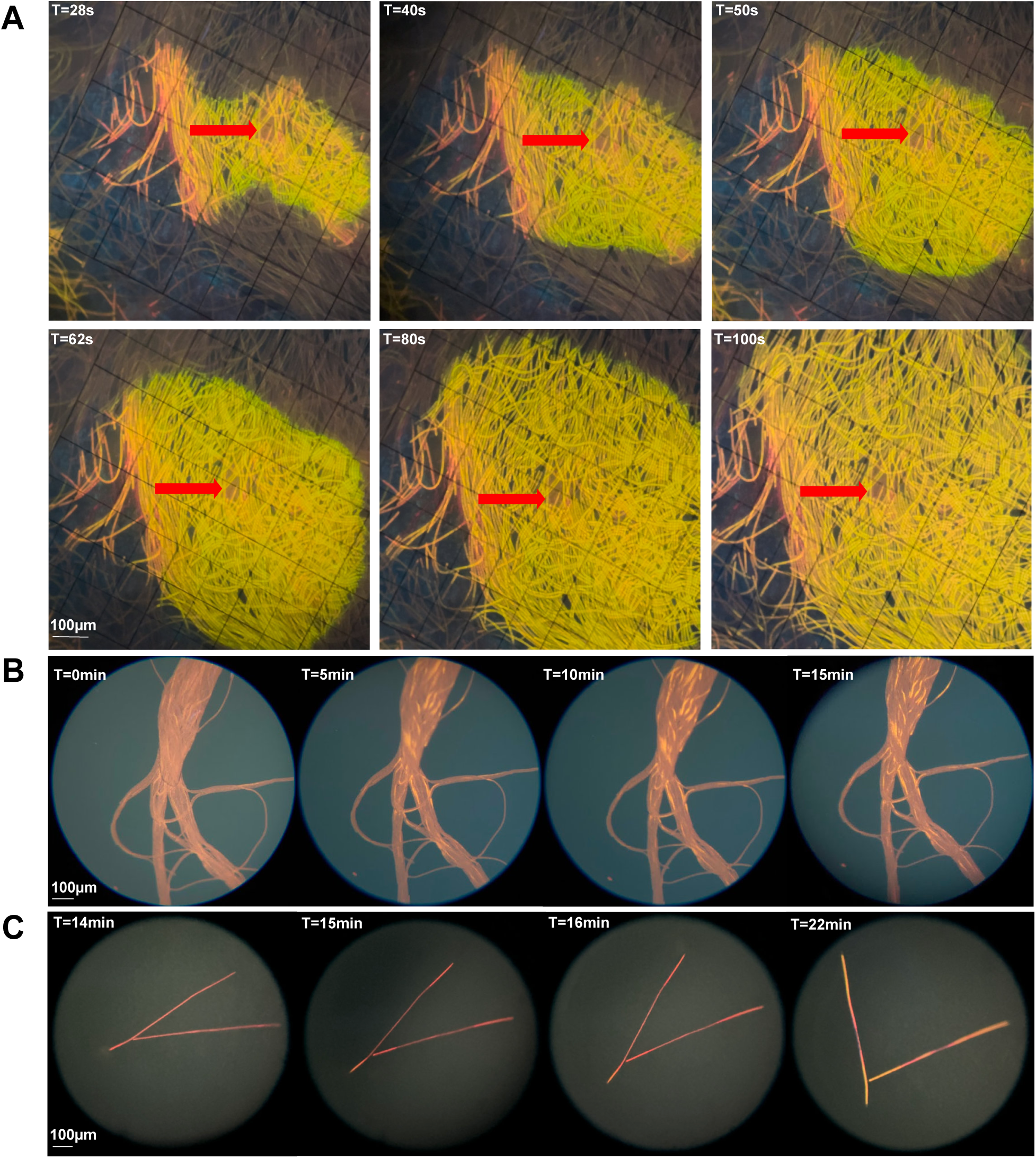
Variability in *Trichodesmium* colony morphology and color-changing response when visualized using DAPI long-pass epifluorescence. (**A**) Wet mounts of the color changing progression of *T. erythraeum* DEMO, T=28.40,50,62,80,100 sec. The red arrow is pointing at the same location on colony at each time point. All 6 pics having the same scale. (**B**) Same processes for *T. erythraeum* GS4, (T=0, 5, 10, 15min), and (**C**) in *T. erythraeum* RLI (T=14, 15, 1, 22min).

Interestingly, this color changing phenotype appeared to be more of a typical TeryB phenomenon as TeryA isolates did not always get brighter, but instead very often they would bleach (i.e., lose its original pigmentation and become pale) with prolonged light exposure (∼5mins) before a color change can appear. For those occasions where TeryA trichomes changed color, the change was found to begin from both ends of the trichome and expand inwards. The whole process would start after ∼10-15mins of light shock and the trichome’s color would change from red to golden over the next ∼5mins (**Fig. 3C**).

We further tested if the color change phenotype is a direct response only activated in light exposed cells but absent in adjacent non-exposed ones. Specifically, colonies of TeryB-2 were exposed to light shock under 40x magnification for 10mins. After the intense light shock, once turned back to 10x magnification, a clear sharp-edge circle shape could be seen which did not spread to adjacent non-shocked cells (**Fig. S1**). These data indicate the color change phenotype is a restricted, fast response of only the cells under strong, blue-shifted light stress (i.e., near UV), and could indicate a phenotypical change of TeryB-2 in response to the intense blue light in oligotrophic habitats.

### Pangenome analysis revealed preserved & unique gene clusters in each subclade

As pangenome analysis includes all genes for a given group of genomes (Eren et al., 2021), the presence/absence of homologous gene clusters (HGC) provides the opportunity to understand genetic variation and conservation in microbial taxa. In our *Trichodesmium* pangenome analysis, we also included three high-quality MAGs representing TeryA-2 and TeryB-1 (Delmont, 2021; Koedooder et al., 2022; Webb et al., 2023), in addition to the isolate genomes, to supplement the genome collection of those lineages (**Fig. 4**). We identified 1730 gene clusters in the single copy core genome (SCG) of *Trichodesmium* conserved by all 14 genomes included in this study (**Fig. 4**), which takes up 30%-36% of all gene clusters in each genome. We also identified 602 extended core gene clusters in the *Trichodesmium* pangenome, which represent gene clusters that are preserved by all genomes but can have multiple copies. The identification of the SCG core is especially critical for further understanding *Trichodesmium* distribution in global ocean samples because it provides a more complete reference for metagenomic read recruitment, excludes the potential biases caused by uneven gene copy numbers in different genomes, and eliminates concerns of taxonomically broadly-conserved selfish DNA elements. Furthermore, we also defined subclade-conserved HGCs (**Fig. 4**) that correlate with their phylogenetic placement and/or relate to their unique phenotypic characteristics of each subclade.

**Figure 4.**
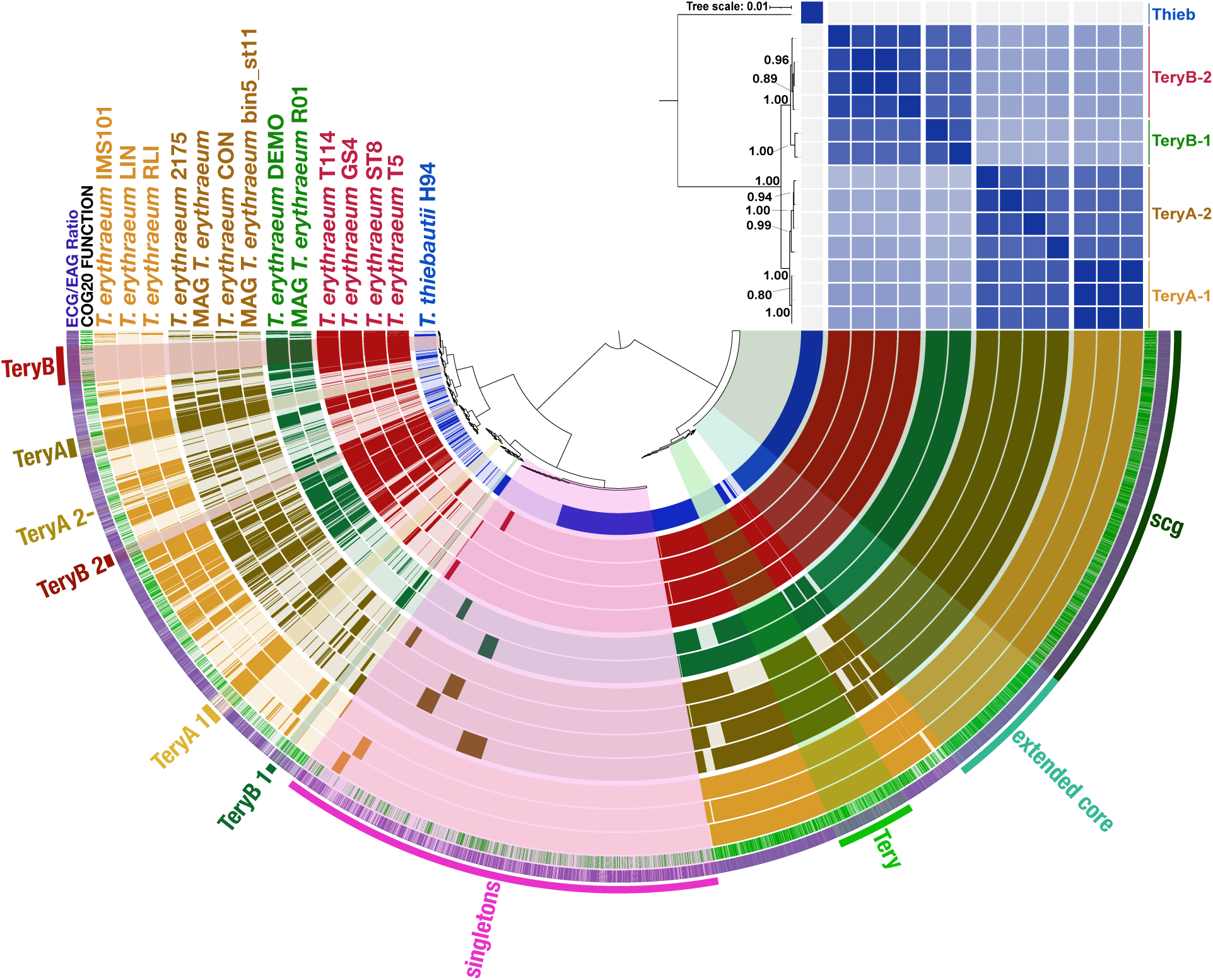
*Trichodesmium* pangenome analysis with 14 total genomes (11 isolate genomes and 3 MAGs) representing *T. thiebautii* and four subclades within *T. erythraeum*. Colored ring on the pan represents (from inner to outer): genomes from Thieb, TeryB-2, TeryB-1, TeryA-2, TeryA-1, COG annotations, and ECG/EAG ratio (i.e., environmental core or accessory gene ratio). Presence and absence of thin bars on each genome layer indicates presence/absence of gene clusters. Presence and absence of thin green bars on the COG ring indicate COG annotated/non-annotated of each gene cluster. Color density of thin purple bar on the ECG/EAG ratio ring indicate detection level of that gene cluster in field samples. Additional layer on the top right is showing the average nucleotide identity of the genomes, with a phylogenetic tree attached to its left (Bootstrap values above 0.5 are shown). Abbreviations: SCG = single copy core gene clusters; singletons = gene clusters only present in one of the genomes; extended core = conserved homologous gene clusters that are present in multiple copies; and colored clade labels on the most outer ring indicates group-specific conserved gene clusters. The pangenome does not show synteny.

#### a. HGC revealed most differences between Thieb and Tery

Like the findings of Webb et al., 2023, the most HGC differences were found between the Thieb and Tery clades. Based on our pangenomic analysis, there are in total of 368 HGC that only exist in the Tery clade. This major difference can also be supported by other evidence: (1) the phylogenetic placement using 251 core genes of cyanobacteria shows the deepest branching separating Thieb H94 with other *Trichodesmium* genomes (**Fig. 1&4**); (2) GC-content of Thieb is higher than any Tery (**Table S1**); (3) whole genome average nucleotide identity (ANI) analysis showed the lowest genome similarity between Thieb and any subclades in Tery (<90%) (**Fig. 4 & Table S2**). However, since we only had one Thieb isolate genome available at the time of this work, herein we will focus on the Tery pangenome and phenotypic trait mapping.

#### b. Genomes of TeryB are genetically different from TeryA

As mentioned, although all members of the Tery clade have very high ANI (>97%), phylogenomic analyses showed strong bootstrap support for the split of TeryA and TeryB subclades (**Fig. 4)**. Corroborating this finding, pangenomics determined that members of TeryA & B each have conserved auxiliary gene clusters absent in the other group (i.e., 87 TeryA and 178 TeryB unique HGC; **Fig. 4**). Of these HGCs, TeryA and TeryB have 26 and 43 annotated with predicted functions, respectively (Table S3). Of these genes, histidine ammonia-lyase (*hutH*), chitinase gene (*chiA*) are discussed more below and they are genes predicted to be involved in important nutrient acquisition steps.

Of all the Tery genomes in this study, only TeryB maintain histidine ammonia-lyase (*hutH*). Studies have shown that histidase (product of gene *hutH*) is involved in the first step of histidine degradation, where an α-amino group is removed from the compound (Gerth et al., 2012; Itoh et al., 2007). Thus, histidine, a common marine molecule that is enriched in carbon and nitrogen (Bender, 2012; Xian et al., 2022), could potentially be an important nutrient resource for TeryB. We did not, however, find expression signals of this gene in *Trichodesmium* metatranscriptomic samples from both the North Atlantic and North Pacific (**Fig. S2**; (Cerdan-Garcia et al., 2022; Frischkorn et al., 2018; Rouco et al., 2018), indicating this gene might undergo selective stress, being suppressed in those areas, or be undetectable due to lower amounts of TeryB present in those samples (**Table S4**).

The chitinase gene *chiA* is predicted to carry out a step in chitin utilization, which could be a potential, important alternative source of organic carbon and nitrogen (Capovilla et al., 2023). Interestingly, the *chiA* gene is conserved in TeryB and Thieb H94, but is absent from all TeryA genomes. Read recruiting RNAseq datasets show that *chiA* from TeryB subclades is expressed *in situ* in RNA samples obtained from both the North Pacific and Atlantic Oceans (**Fig. S2**). The expression data suggests that TeryB members could be actively augmenting their C&N pools with chitin. Contrastingly, it is surprising that both TeryA subclades are missing this alternative acquisition gene, especially when one considers their habitat is the oligotrophic ocean. Previous studies with the marine cyanobacteria *Prochlorococcus* showed that, for this planktonic genus, there’s a relationship between the absence of chitin acquisition genes and genomic streamlining (Capovilla et al., 2023). With limited *Trichodesmium* genome numbers, it is difficult to determine if TeryA subclades are indeed under the process of genome streamlining (as described in (Giovannoni et al., 2014)), however, their genomes are lower in GC percentage compared to other subclades, which is also one of the indicators of genome streamlining processes (Giovannoni et al., 2014).

#### c. Subclades within TeryA/B can have different genetic potential

We found annotated functional genes that could contribute to the difference and similarity between TeryA-1 and TeryA-2. There are total of 110 gene clusters different between the two TeryA subclades (**Fig. 4**, **Table S3**). While many of these gene clusters are not annotated (70/110) or not showing an obvious subclade pattern relevant to discuss here, the photosystem I related gene PsaM stood out as being of potential importance.

All members of TeryA-1 are missing the gene predicted to encode Photosystem I (PSI) reaction center (subunit VII, PsaM). Based on the literature, while other cyanobacterial strains defective in this gene do not have an observable growth or photosynthetic defect (Fromme et al., 2001; Naithani et al., 2000), it is likely that the absence of this gene is still impacting TeryA-1 strains ability to form a stable PSI trimer and result in less chlorophyll content per cell (Naithani et al., 2000). This could mean that members of the Tery-A1 might be more adapted to higher lights, while TeryA-2 group are better at photoacclimation to lower light intensities.

The pangenome also identified a group of HGCs only conserved in TeryB-2 but missing in TeryB-1 (55 HGCs; **Fig. 4**). More than 30% of these gene clusters (19/55) were annotated with functions. They were mostly enriched in amino acid/coenzyme/lipid/secondary metabolites transport and metabolism (**Table S3**). Hypothetically, some of these gene clusters could be contributing to the color changing phenotype of the TeryB-1 DEMO, but we could not identify an annotated functional gene implicated in this phototrophic process (**Table S2**). While it is possible that the genes involved in DEMO’s unique color changing phenotype are unannotated, this absence combined with the speed of the change suggests a potential role for post-translational modification and thus calls for future proteomic analysis in this subclade.

#### d. Specific subclades maintained different paralog copies of predicted Na+/proline symporter

Gene copy numbers can have unexpected phenotypic impacts due to dosage effects (Schirrmeister et al., 2012), and we discovered that the Tery subclades have different copy numbers for many predicted genes. For example, TeryA-1 and TeryA-2 isolates all encode 5 copies of a predicted Na+/proline symporter (PutP), while all TeryB isolates have 6 and Thieb H94 has 8 copies. Homologs of this symporter have been documented as one of the pathways that marine bacteria can use for adapting to higher salinity (Dufresne et al., 2003; Jung et al., 2012). Less copies could be disadvantageous for TeryA-1 and TeryA-2 in saltier water. Fittingly, the expression of this gene can be seen in both oceanic Atlantic and Pacific samples (**Fig. S2**).

### Global distribution and in situ activity of *Trichodesmium* clades

We used the conserved single copy core genome in each clade defined in our pangenomics workflow (i.e., TeryA, TeryB, Thieb) to reveal *Trichodesmium* biogeography and activity in bioGEOTRACES, GO-SHIP, station ALOHA and BATS bulk water sample datasets (Biller et al., 2018; Larkin et al., 2021). Because of high intra-clade ANI of members of TeryA and B (**Fig 4**), we were not able to further perform biogeography down to the subclade level (e.g., Tery A1 & 2 etc). However, the whole-water metagenomic results revealed an uneven TeryA, TeryB, and Thieb distribution pattern in global basins with Thieb significantly more abundant in most oceans (**Fig. 5A**). Interestingly, this read mapping also showed that TeryA and TeryB relative abundance is consistently higher in the Bay of Bengal (**Fig. 5A inset**), with more TeryA in the north and relatively more TeryB in the south. We compared the metadata of sampling locations and found an increasing representation of Tery in samples above 29°C (**Fig. S3**) supporting their predicted better adaptation to warmer environments compared to Thieb (Chappell & Webb, 2010). But in most of those warmer samples (**Fig. 5A lower inset**), Thieb still dominates the relative abundances, indicating some other environmental factor(s) in the Bay of Bengal (BOB) are responsible for the reversed pattern where increased Tery are observed.

**Figure 5.**
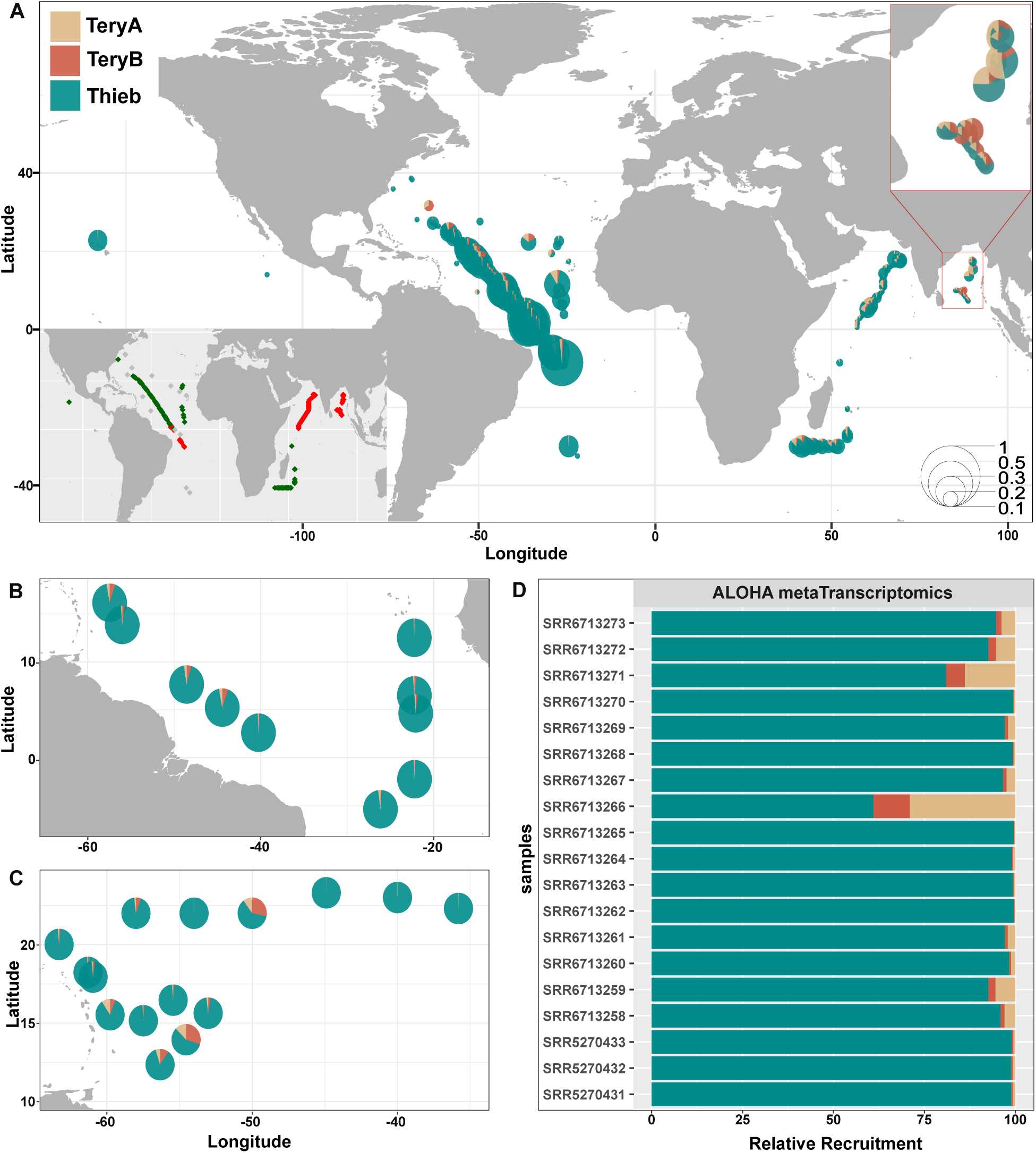
Metagenomic and metatranscriptomics readmapping revealed *Trichodesmium* subclades global distribution pattern. (**A**) Metagenomic readmapping using bulk water samples from BioGEOTRACES, GO-SHIP, station ALOHA and BATS. Diameter of the pie indicates total *Trichodesmium* abundance in that environmental sample (scale legend on lower right corner, unit: square root % of mean coverage values). Portion of each pie slice shows the relative abundance of each subclade. Upper right corner shows a zoom-in view of samples in the Bay of Bengal (the non-magnified pie sizes apply to the scale legend). Lower left corner shows a color-coded view of temperature log at most of the sampling stations (Red: >=29°C Green: <29°C). (**B**) *Trichodesmium* subclade distribution pattern from hand-picked *Trichodesmium* colony metagenomes in a North Atlantic Transect (TriCoLim, (Webb et al., 2023)). Pie diameter set to r=1, portion of each pie slice shows relative abundance of each species. (**C**) Relative gene expression activities of *Trichodesmium* subclades in North Atlantic using transcriptomic samples from previous studies (Cerdan-Garcia et al., 2022; Rouco et al., 2018). Pie diameter set to r=1, portion of each pie slice shows relative representative of each species. (**D**) Relative gene expression activities of *Trichodesmium* subclades at station ALOHA, Pacific using metatranscriptomic samples (Frischkorn et al., 2018; Rouco et al., 2018)

Based on the genomic potential discussed above, we hypothesize that this BOB Tery enrichment factor could be salinity. A calculated 18-year average salinity dataset (2001-2018) showed that the BOB’s salinity varied between 31 and 34, significantly lower than the other oceans, including the Arabian Sea at the same latitude on the other side of the Indian peninsula (Li et al., 2021). Data from the GO-SHIP cruises (I09 and I07) corroborated this and showed that BOB salinity (Cruise I09), where we found the highest Tery relative abundances, is comparatively lower than the rest of the ocean (< 33 PSU; **Fig. S4**). Specifically, North of 5°N in I07 and I09, mean surface salinity was significantly higher in the Arabian Sea (35.76 ± 0.55 PSS-78, *n* = 35) than in the Bay of Bengal (32.96 ± 0.59 PSS-78, *n* = 44). The difference was highly significant using both Welch’s two-sample *t*-test (*t* = 21.77, *p* < 10⁻³³) and the nonparametric Mann–Whitney U test (*U* = 1540, *p* < 10⁻¹³). Fittingly, most of our new Tery isolates in this study came from media made with 75% Sargasso Sea water (i.e., sanity ∼26 PSU). Additionally, our data show that TeryA is relatively more abundant in the inner BOB (**Fig. 5A upper inset)**, implying that temperature and salinity could act in concert to select for TeryB in the fresher, warmer waters east of Sri Lanka. While these data support the role of salinity in Tery relative abundances in general, more research is required to discern the differences observed between Tery A and B in situ.

These data add a nuanced wrinkle to the hypothesis that Tery abundances will increase due to climate change globally. For example, previous studies have indicated that TeryA isolates grow faster than Thieb and have higher nitrogen fixation rates under variant temperature and Fe availabilities (Chappell & Webb, 2010), highlighting the potential important nitrogen contribution of this subclade in the warmer oligotrophic oceans. Physiological experiments of TeryA also showed its better growth and higher nitrogen fixation rate under increasing temperature and [CO_2_] (Barcelos e Ramos et al., 2007; Boatman et al., 2020; Hutchins et al., 2007, 2013), and thus were hypothesized to be more successful under global climate change conditions. Our data show the relative abundance of TeryA/B is correlated with likely lower salinities. Experimental studies should rigorously probe the salinity requirements of *Trichodesmium* subclades to confirm whether they are important drivers of abundance and physiology.

To understand genomic composition within *Trichodesmium* colonies, we also used metagenomic reads from hand-picked North and Equatorial Atlantic *Trichodesmium* colonies (Webb et al., 2023). Read mapping result showed similar patterns to what we have seen in bulk water samples: Thieb numerically dominates this oceanic region, with only limited observation of TeryA and TeryB (**Fig. 5B**). The corroborating results from multiple sources of bulk water metagenomic samples also relieves concerns that hand-picking colonies would be biased against TeryA due to their frequent free-trichome lifestyle (e.g., Hynes et al., 2012).

Furthermore, using metatranscriptomic reads from hand-picked colonies (Cerdan-Garcia et al., 2022; Frischkorn et al., 2018; Rouco et al., 2018), we revealed the relative gene expression activity of *Trichodesmium* clades aligns with their relative numerical abundance pattern in the North Pacific and North Atlantic (**Fig. 5C & D**). One interesting thing to notice is that although Tery is still less frequently detected than Thieb, we see a higher variation in gene expression level of TeryA and TeryB, data that suggest a potential opportunist’s role in fluctuating environmental conditions.

## Conclusion

Here we present new high-quality genomes of *T. erythraeum* representing two major clades (TeryA and B) that can further be broken down into four subclades (TeryA-1&2, and TeryB-1&2) based on phenotype variation, ANI, and genomic content. Despite the fact that all previous *T. erythraeum* strains have been enriched in phycoerythrobilin and were reddish in color (Hynes et al., 2012), our new mid-Atlantic isolate, TeryB-1 DEMO, grows as green-colored colonies in higher light. In combination with data from the extinct Thieb H94 that was also greenish in color, our findings indicate that green *Trichodesmium* isn’t always dead as previously argued (Neveux & Tenório, 2008; Orcutt et al., 2008). On the contrary, our data suggest that green *T. erythraeum* DEMO appear to be better adapted to blue light than all previously characterized Tery members (Hynes et al., 2012). Additionally, this study highlights that all Tery genomes are very closely related and shows that the classical species names for the genus based on morphology are difficult to reconcile with genomics and physiology. Thus, we argue that the most accurate way to distinguish different *Trichodesmium* spp. is using isolate pangenomics and cladistics.

We show that Tery and Thieb global distributions are uneven with the latter relatively dominant in whole water column and picked colony datasets. The clade that we know the most of about (i.e., many controlled physiology and evolutionary datasets) is TeryA-1 (*i.e., T. erythraeum* IMS101), the least commonly observed, major clade of *Trichodesmium* in situ. This calls for more field investigations and additional, in-depth physiological experiments with Thieb cultures to better predict the clades abundance and activity in the future ocean.

Finally, while it is predicted that increasing CO_2_ concentrations and temperatures caused by climate change will likely benefit TeryA-1&2 growth more than other *Trichodesmium* subclades (Hutchins et al., 2013), our discovery of Tery enrichment in lower salinity regions like the BOB may weaken this prediction. This is because the future oligotrophic ocean is forecasted to face more intense vertical stratification causing ocean surface salinity to be more stratified than it is now. This is predicted to result in salty regions getting saltier while fresh regions get fresher (Cheng et al., 2020; Durack et al., 2012). While there may be other factors associated with the observed enrichment of Tery in the BOB, our results suggest that ocean salinity could also have a major influence on their distribution and activity.

## Acknowledgements

We acknowledge the captain and crew of the R.V. Atlantis for their assistance in acquiring the new USC cultures and John B. Waterbury for providing WHOI *Trichodesmium* isolates. This work was supported by National Science Foundation grants OCE1657755, OCE185122, BIO2125191. Reads are available at NCBI SRA under accession number PRJNA1460435.

