## Supplemental Figures for "Global distribution of cultivated *Trichodesmium* subclades revealed by multi-omics"

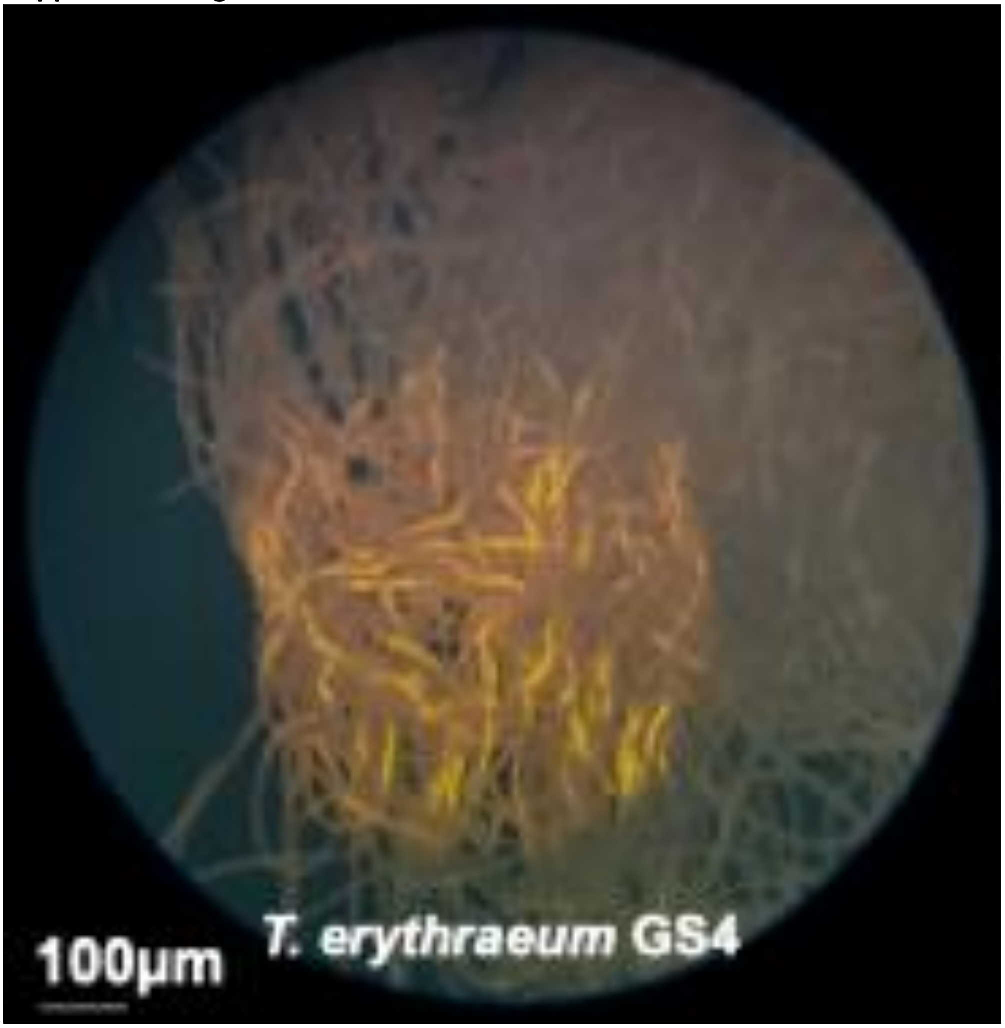


Supplemental Figure 1. ‘Lighting-up’ response of TeryB-2 GS4. Lighted circle in the figure showing the region that was exposed to mild UV light stress for 10mins.

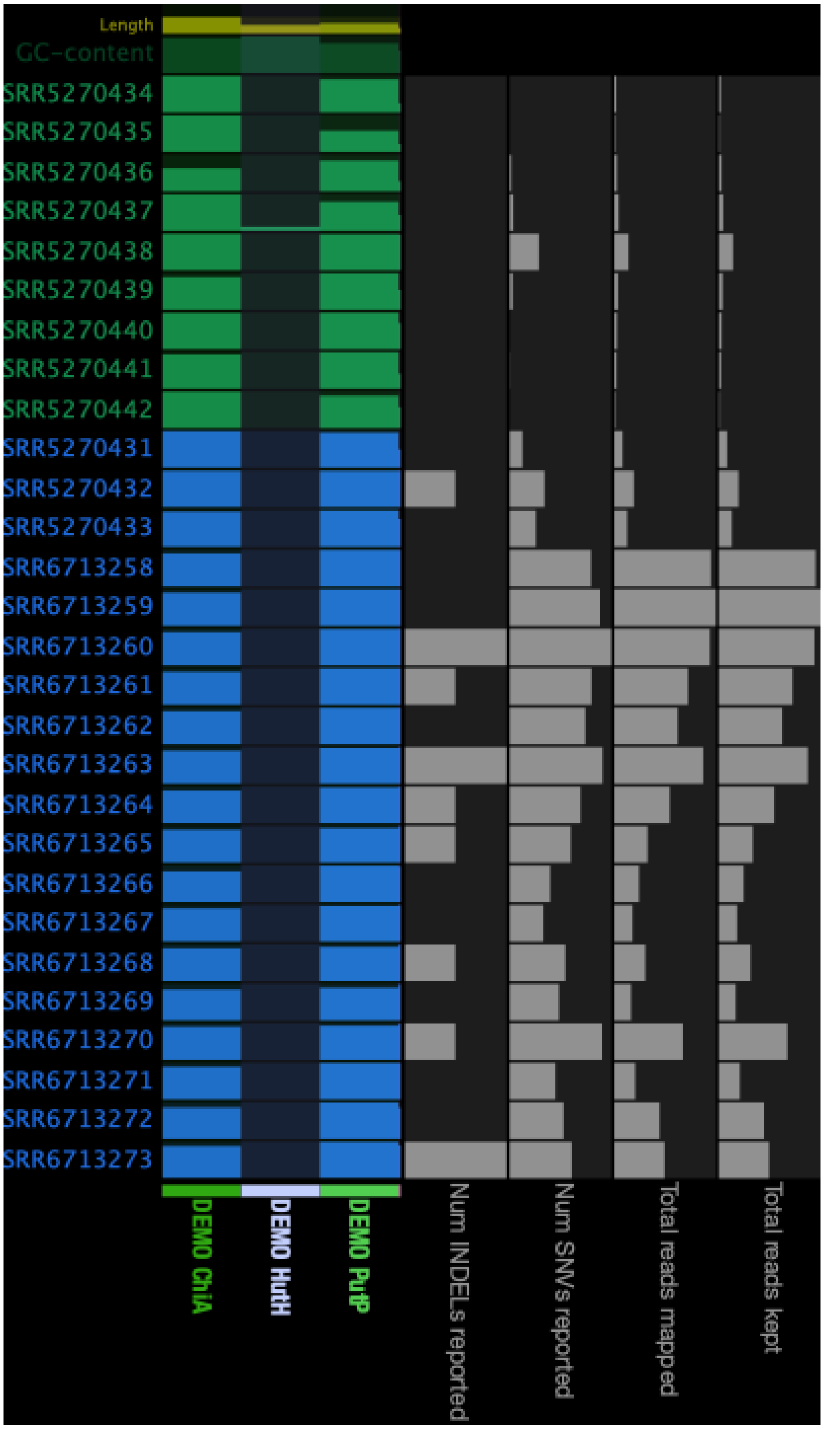


Supplemental Figure 2. Expression pattern of Tery subclades differentially preserved genes in both North Atlantic (green SRR#s) and North Pacific (blue SRR#s). Strain names prior to gene name indicates the source of the specific sequence used for read-mapping.

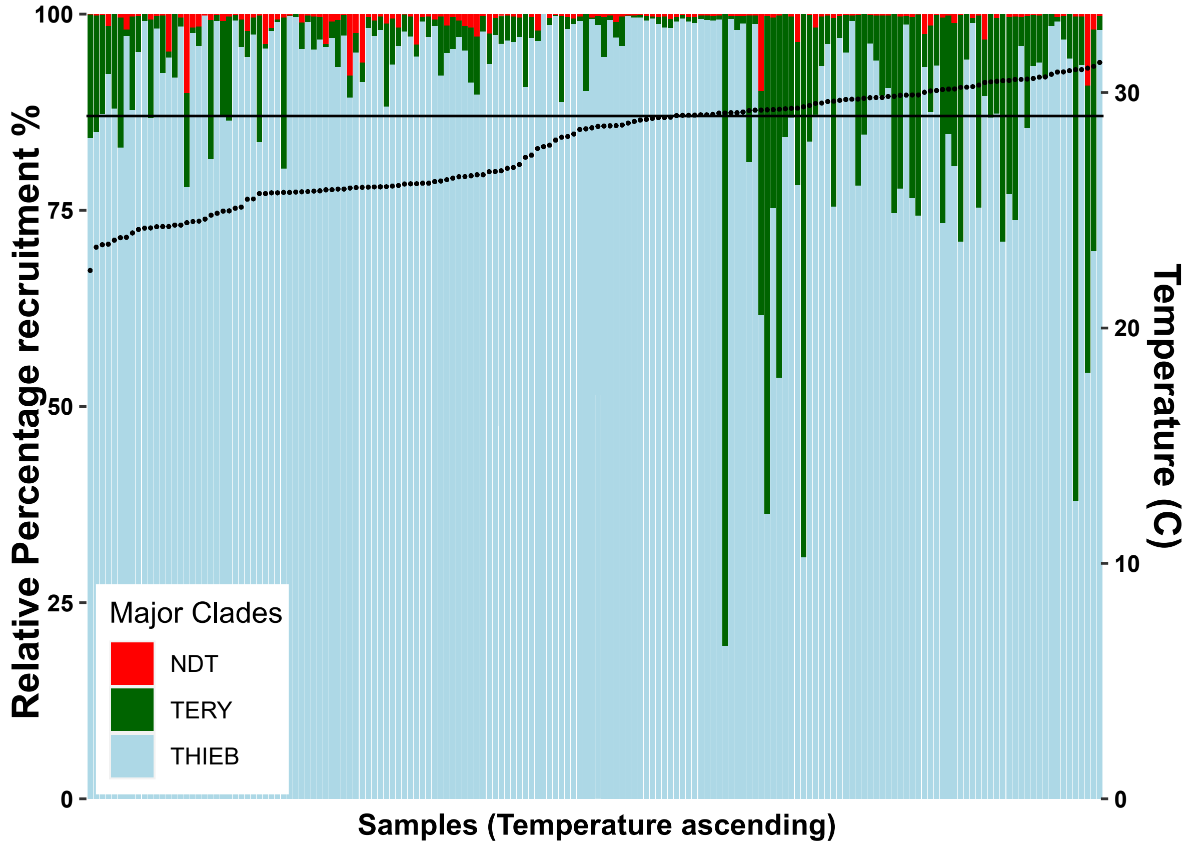


Supplemental Figure 3. Relative abundance of *Trichodesmium* major two clades and newly found Non-diazotrophic *Trichodesmium* (NDT) in bulk water samples with increasing temperature. Dot in each sample column indicate the temperature mark and the horizontal line represents the threshold of 29°C.

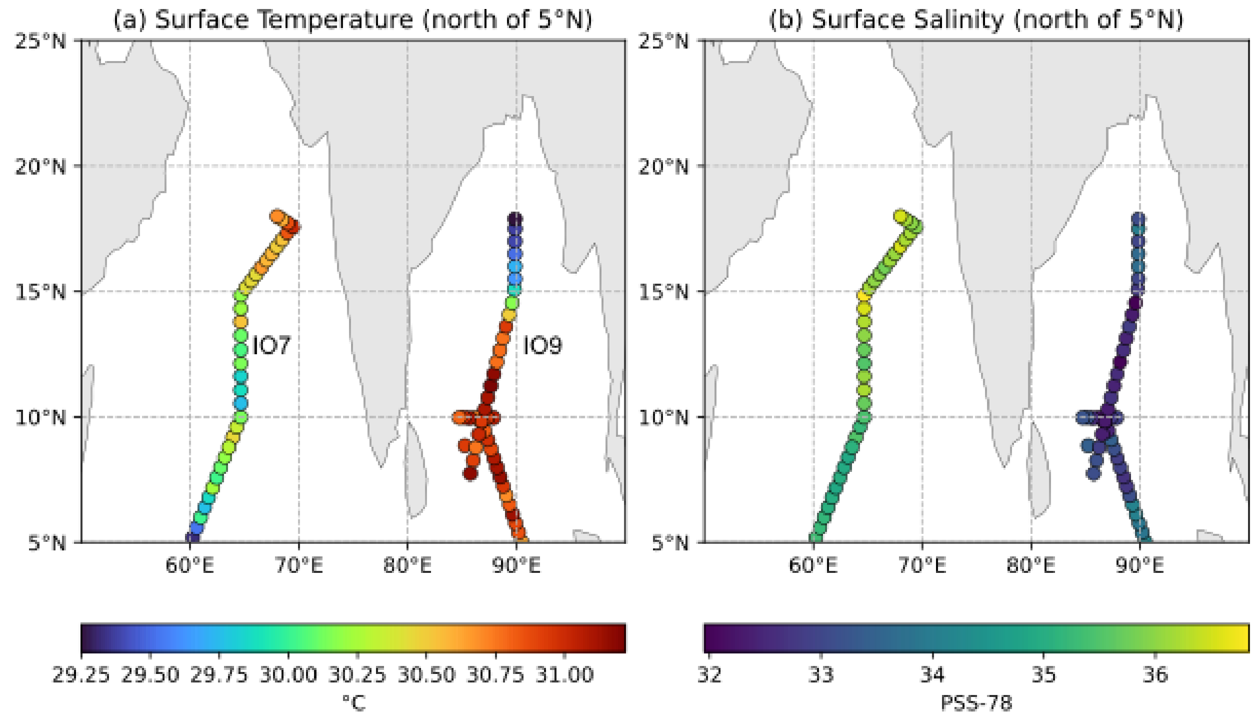


Supplemental Figure 4. Surface salinity and termperature of GOSHIP cruise I07N 2018 and I09N 2016 stations north of 5N of in Arabian Sea and Bay of Bengal, repsectively.
